# Assessing nonlinearities in the GFP random mutagenesis landscape using the Power Transform

**DOI:** 10.64898/2025.12.09.693289

**Authors:** Dmitry A. Petrov, Dmitry N. Ivankov

## Abstract

Epistasis, a non-additive contribution of mutations to fitness, complicates genotype-to-phenotype prediction and is often confounded by nonlinearities in phenotype measurements. The Power Transform method, particularly Box-Cox, has been used to reduce epistasis in small, combinatorially complete datasets by linearizing phenotypic scales. However, its applicability to large landscapes, especially those generated by (quasi-)random mutagenesis, remains unclear. Here, we apply both Box-Cox and Yeo-Johnson Power Transforms to the extensively characterized green fluorescent protein (GFP) fitness landscape, which contains hundreds of thousands of two- and three-dimensional combinatorially complete datasets. Surprisingly, when applied to individual hypercubes, Box-Cox reduces pairwise epistasis by 19.4% on average, whereas Yeo-Johnson – despite accommodating negative values – slightly increases epistasis (by 4.45%). Moreover, in a notable fraction of hypercubes, both methods increase the magnitude of epistatic coefficients, indicating that Power Transform does not universally reduce nonlinearity at the local scale. In contrast, applying Yeo-Johnson to the largest connected component of the GFP landscape (20,872 genotypes) successfully reduces both pairwise (by 3.86%) and third-order epistasis (by 11.6%), with the majority of coefficients decreasing in magnitude. These results demonstrate that Power Transform can be extended to large, real-world landscapes generated by random mutagenesis, but only when applied to connected sublandscapes. Our findings highlight a critical distinction between global and local linearization and caution against assuming that Power Transform always diminishes epistasis.

## Introduction

Epistasis, the dependence of a mutation’s effect on its genetic background, represents a major obstacle to predicting fitness and phenotype from genotype (Cordell, 2002; Phillips, 2008; Moore, Williams, 2009; Poelwijk *et al*., 2009; Verhoeven *et al*., 2010; de Visser *et al*., 2011; Weinreich *et al*., 2013; Ivankov *et al*., 2014; MacKay, 2014; Sailer, Harms, 2017; Zhou *et al*., 2022; Johnson *et al*., 2023). Therefore, uncovering the patterns that govern epistasis is essential for achieving accurate phenotype and fitness predictions. Experimental data to investigate such patterns can be obtained by measuring how fitness and/or phenotype change in response to specific genetic alterations (Poelwijk *et al*., 2009; de Visser *et al*., 2009; Hall *et al*., 2010; Khan *et al*., 2011; Olson *et al*., 2014; Anderson *et al*., 2015; Sarkisyan *et al*., 2016; Pokusaeva *et al*., 2019; Johnston *et al*., 2024). Because fitness is often more challenging to measure than phenotype, researchers frequently use phenotypic measurements as a proxy for fitness (Olson *et al*., 2014; Sarkisyan *et al*., 2016). In this work, we treat fitness as a specific type of phenotype and use the terms “fitness” and “phenotype” interchangeably.

Epistatic coefficients can be calculated from so-called combinatorially complete datasets (Weinreich *et al*., 2013). In such datasets, for a chosen set of *L* sites (e.g., protein positions), each site has two possible alleles (e.g., wild-type or mutant), and all 2^*L*^ possible genotypes have been experimentally assayed. If we assign a binary axis to each mutation (0 – no mutation, 1 – mutation present), this dataset forms an *L*-dimensional hypercube in sequence space (in this work, we use terms “combinatorially complete dataset” and “hypercube” interchangeably). Several studies have deliberately generated such combinatorially complete datasets (Poelwijk *et al*., 2009; de Visser *et al*., 2009; Hall *et al*., 2010; Khan *et al*., 2011; Olson *et al*., 2014; Anderson *et al*., 2015; Johnston *et al*., 2024). Alternatively, mutant libraries can be created using (quasi-)random mutagenesis approaches (Sarkisyan *et al*., 2016; Pokusaeva *et al*., 2019). In the resulting phenotype-genotype maps, epistatic coefficients can still be computed provided one identifies embedded combinatorially complete datasets. All such datasets can be efficiently found using the HypercubeME algorithm (Esteban *et al*., 2019).

The standard methods for quantifying epistasis assume that the phenotypic effects of individual mutations combine additively (Fisher, 1918; Cheverud, Routman, 1995; Phillips, 2008; Weinreich *et al*., 2013; MacKay, 2014; Poelwijk *et al*., 2016). However, if mutations instead interact multiplicatively – or through any other nonlinear relationship – the additive model will infer apparent epistasis even in the absence of true interaction between mutations (Sailer, Harms, 2017). Consequently, any nonlinear transformation of the phenotype will alter the amount of detected epistasis. When analyzing empirical landscapes, it remains unclear how much of the observed epistasis stems from such nonlinearity versus genuine interactions between mutations. This raises a critical question for each landscape under study: what amount of epistasis would be detected if we minimize, as far as possible, the contribution from nonlinearity?

In 2017, Sailer and Harms utilized the Power Transform method with the Box-Cox transformation, to remove nonlinearities from epistatic landscapes (Sailer, Harms, 2017). They analyzed seven small combinatorially complete datasets, each comprising five to six loci (i.e., consisting of 2^5^ = 32 or 2^6^ = 64 genotypes) (de Visser *et al*., 2009; Hall *et al*., 2010; Khan *et al*., 2011; Weinreich *et al*., 2013; Anderson *et al*., 2015). Sailer and Harms clearly demonstrated that five of these seven datasets still exhibited higher-order epistasis even after applying the Power Transform to correct for nonlinearity (Sailer, Harms, 2017). Thus, applying the Power Transform to individual combinatorially complete datasets offers significant promise for reducing epistasis caused by nonlinear scaling. The next logical step is to extend this approach to entire fitness landscapes – particularly those generated by (quasi-)random mutagenesis (Sarkisyan *et al*., 2016; Pokusaeva *et al*., 2019).

In this study, we explore the application of the Power Transform to an entire landscape using the example of the classic landscape of green fluorescence protein (GFP) obtained via random mutagenesis (Sarkisyan *et al*., 2016), which contains numerous two- and three-dimensional hypercubes (i.e., combinatorially complete datasets of size 2^2^ = 4 and 2^3^ = 8). When we apply the Box-Cox-based (Box, Cox, 1964) Power Transform (as in Sailer, Harms, 2017) to individual hypercubes, pairwise epistatic coefficients decrease on average by 19.4%, and third-order coefficients decrease by 1.30%. Surprisingly, however, when we use the more flexible Yeo-Johnson transformation instead, epistatic coefficients *increase* on average – by 4.45% for pairwise epistasis and by 0.43% for third-order epistasis.

Applying the Power Transform to the entire landscape proved feasible only when the landscape is connected. Using the Yeo-Johnson (Yeo, Johnson, 2000) transformation (the Box-Cox transformation failed due to its restrictions), we applied the Power Transform to the largest connected sub-landscape of GFP, which comprises 20,872 genotypes. In this case, the average pairwise epistatic coefficient decreased by 3.86%, and the third-order coefficient decreased by 11.5%.

Our results demonstrate the presence of higher-order epistasis in the GFP landscape both before and after the application of the Power Transform method, supporting earlier conclusions about the importance of higher-order epistasis in empirical fitness landscapes (Weinreich *et al*., 2013; Sailer, Harms, 2017). Our findings also reveal that the effectiveness of nonlinearity removal in individual hypercubes depends critically on the choice of transformation – Box-Cox (Box, Cox, 1964) versus Yeo-Johnson (Yeo, Johnson, 2000). Most importantly, we demonstrate that the Yeo-Johnson-based Power Transform can be applied to any large landscape – provided it is connected.

## Materials and Methods

### Identification of combinatorially complete datasets

In the GFP landscape (Sarkisyan *et al*., 2016), we found 408,673 two-dimensional and 1,446 three-dimensional combinatorially complete datasets using the HypercubeME algorithm (Esteban *et al*., 2019).

### Power Transform with Box-Cox transformation

The approach considered by Sailer and Harms to reduce nonlinearities in fitness landscapes (Sailer, Harms, 2017) uses the Power Transform method with a modification of the Box-Cox power transformation (Box, Cox, 1964). For each single mutation present in the landscape, the averaged effect across all genetic contexts is computed, and then, based on these effects, an additive phenotype is calculated for each genotype (Sailer, Harms, 2017). The stepwise approach is as follows (Sailer, Harms, 2017):

1. Compute approximations of additive phenotypes:

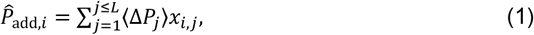

where ⟨Δ*P*_*j*_⟩ is the average effect of mutation *j* across all backgrounds calculated from the measured phenotypes *P*_obs_, *x*_*ij*_ is an index that encodes whether or not mutation *j* is present in genotype *i*, and *L* is the number of sites.
2. Fit a nonlinear least squares regression of observed phenotypes vs. approximations of additive phenotypes:

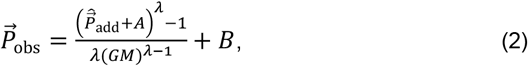

where *A* and *B* are translation constants, *GM* is the geometric mean of 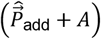, and *λ* is a scaling parameter.
3. Apply the inverse transformation to the observed phenotypes using the fitted parameters, thereby linearizing observed phenotypes:

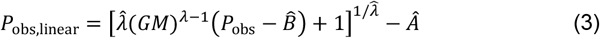

When we attempted to use the original code from (Sailer, Harms, 2017) on test combinatorially complete datasets of 4 and 8 genotypes, the program crashed in more than half of the cases. To investigate this issue, we implemented our own version following the equations in (Sailer, Harms, 2017). We wrote the implementation in Python 3, using the NumPy, SciPy, and scikit-learn libraries.

Specifically, we first computed additive phenotypes as defined in Eq. 1. Then, to obtain an initial guess for the Box-Cox parameter *λ*, we applied the power_transform function from scikit-learn library to the observed versus additive phenotype data. Using this initial guess, we performed a refined fit of the nonlinear model (Eq. 2) using scipy.optimize.curve_fit function. Our implementation succeeded in a higher fraction of cases than the implementation from (Sailer, Harms, 2017). Therefore, we used our implementation for all subsequent analyses involving the Box-Cox-based Power Transform.

All Python scripts are available at https://github.com/ivankovlab/power_transform.

### Power Transform with Yeo-Johnson transformation

Note that the Box-Cox-based (Box, Cox, 1964) Power Transform cannot be applied when either the observed or additive phenotypes take non-positive values. Although measured phenotypes in the GFP landscape are strictly positive, computed additive phenotypes can occasionally become zero or negative during analysis, which makes the Box-Cox transformation (Box, Cox, 1964) inapplicable. To address this limitation, we adopted an alternative formulation of the Power Transform based on the Yeo-Johnson transformation (Yeo, Johnson, 2000), which can accommodate non-positive values:

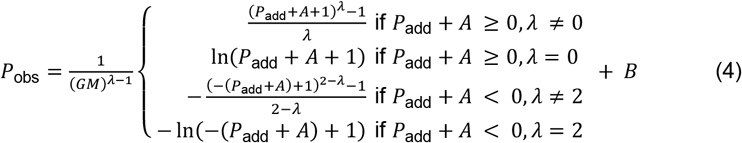

We fit this model to our data; the inverse transformation looks as follows:

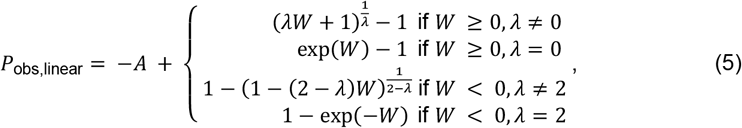

where *W* = (*P*_obs_ − *B*)(*GM*)^*λ*-l^.

### Set of combinatorially complete datasets

To enable a direct comparison between the Box-Cox (Box, Cox, 1964) and Yeo-Johnson (Yeo, Johnson, 2000) transformations, we restricted our analysis to hypercubes for which *both* methods succeeded (i.e., produced valid results without errors). Of the 408,673 two-dimensional hypercubes, both transformations succeeded for 403,162 hypercubes. For all 1,446 three-dimensional hypercubes, both methods succeeded without failure.

### Identification of connected components

We identified connected components of the landscape in two stages: (1) constructing from genotypes with measured phenotypes a connectivity graph in sequence space and (2) detecting connected components in the graph. We represented each genotype as a node and connected two nodes by an edge if Hamming distance between them equaled 1 (i.e., genotypes differed at exactly one amino acid position).

To avoid the O(*N* ^2^) complexity of pairwise comparison (where *N* is the number of genotypes), we generated instead all possible single-mutation neighbors for each genotype – considering the twenty standard amino acids plus the position encoded by a stop codon (overall 21 symbols) – and checked whether each neighbor existed in the landscape. This changed the complexity to *O*(*N*⋅*M*⋅*K*), where *M* is the number of variable positions and *K* = 21. We then applied a standard breadth-first search algorithm to identify connected components (Bundy, Wallen, 1984).

We found that the experimental GFP landscape contains only one large connected component with more than two genotypes, comprising 20,872 genotypes.

### Calculation of epistatic coefficients

We calculated epistatic coefficients according to the Walsh-Hadamard matrix (Weinreich *et al*., 2013; Poelwijk *et al*., 2016):

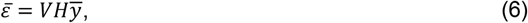

where 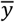 is the vector of phenotypes within a combinatorially complete landscape, 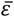 is the vector of background-averaged epistatic terms; *H* is the operator for background-averaged epistasis, defined recursively as:

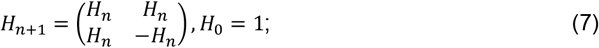

the recursive definition for the weighting matrix *V* is

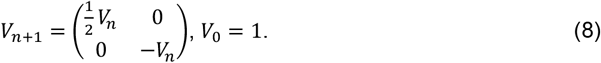

Following Weinreich *et al*. (2013), we quantified the total amount of epistasis as:

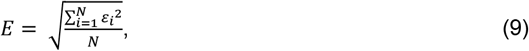

where *ε* are the epistatic coefficients of the highest order present in each hypercube, and *N* is the number of the hypercubes embedded in the full landscape.

### Statistical comparison of distributions

To assess whether Power Transform significantly altered distribution of epistatic coefficients, we performed nonparametric statistical tests. We used two-sided Mann-Whitney U test to compare the full distributions of epistatic coefficients before and after transformation, and one-sided Mann-Whitney U test to compare the distributions of their absolute values.

### Estimating uncertainty in epistatic coefficients

We did not find a way to incorporate measurement error in the Power Transform method. Therefore, to estimate the uncertainty in average value of epistatic coefficients resulting from measurement error in GFP fluorescence, we performed ten runs of the Power Transform for each transformation (both Box-Cox and Yeo-Johnson), using phenotype values randomly generated according to their experimentally determined value and standard deviation. As the standard deviation for any GFP variant, we took value of ± 0.1 estimated from 3642 wild-type GFP measurements (Sarkisyan *et al*., 2016). For each epistatic coefficient, we calculated its standard deviation across these runs, took the average of these standard deviations over all epistatic coefficients 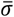, and divided it by the square root of the number *n* of epistatic coefficients to get the estimate of uncertainty *σ*_*x*_ in average value of epistatic coefficients:

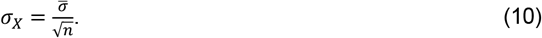

## Results and Discussion

### Dataset

To investigate and illustrate the capabilities and potential of the Power Transform for experimental fitness landscapes, especially those generated by (quasi-)random mutagenesis, we focused on a single, yet extensively characterized, classic landscape of green fluorescence protein (GFP) (Sarkisyan *et al*., 2016). That study resulted in 1,817 single mutations across 233 positions in GFP, either individually or in combinations (up to mutants carrying fifteen single mutations), yielding fluorescence measurements for 51,715 missense variants from the neighborhood of wild-type GFP (Sarkisyan *et al*., 2016).

The GFP landscape is nonlinear; its nonlinearity is largely of a “threshold” nature (Tawfik, 2010; Sarkisyan *et al*., 2016). Protein stability is the most likely factor underlying both this nonlinearity and the overwhelming majority of observed uni-dimensional epistasis.

The GFP landscape contains 408,673 two-dimensional hypercubes and 1,446 three-dimensional hypercubes (see Materials and Methods). The availability of three-dimensional hypercubes is particularly valuable, as it enables robust detection of higher-order epistasis – in our case, third-order epistasis.

### Application of Power Transform to individual combinatorially complete datasets

To assess how the amount of epistasis changes after applying the Power Transform, we computed epistatic coefficients before any transformation and after applying the Power Transform with either Box-Cox or Yeo-Johnson variant. It should be noted that, among all 408,673 two-dimensional hypercubes, both the Box-Cox and Yeo-Johnson versions succeeded for a common subset of 403,162 hypercubes; therefore, we report all subsequent calculations and results only for this group.

Before any transformation of the phenotypes, the average epistasis (calculated as the root-mean-square of all epistatic coefficients) was 0.6741 ± 0.0002 for pairwise epistasis and 0.460 ± 0.003 for third-order epistasis. The corresponding distributions are shown in Figure 1A (pairwise) and Figure 2A (third-order).

**Figure 1.**
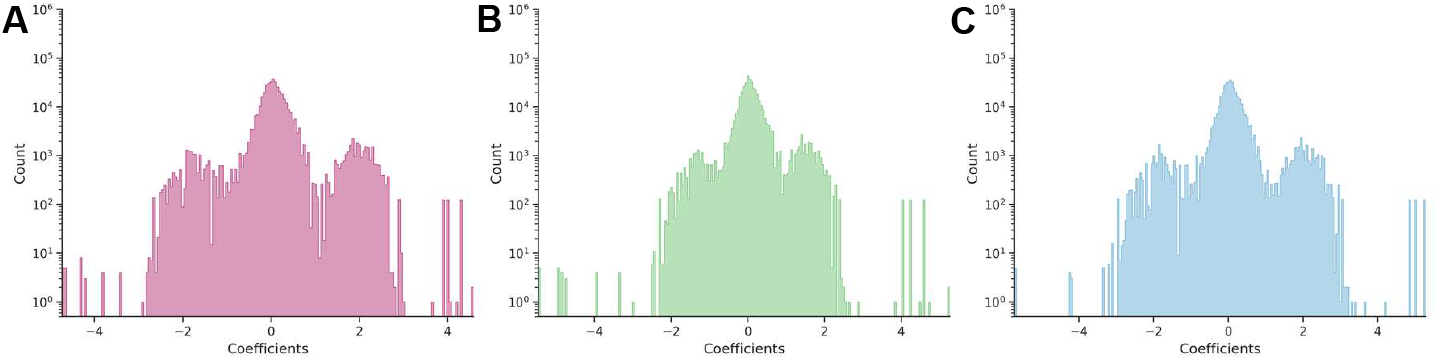
Distribution of pairwise epistatic coefficients computed (A) before Power Transform, (B) after the Box-Cox-based Power Transform, and (C) after the Yeo-Johnson-based Power Transform. In both (B) and (C) cases, the Power Transform was applied separately to individual two-dimensional combinatorially complete datasets.

**Figure 2.**
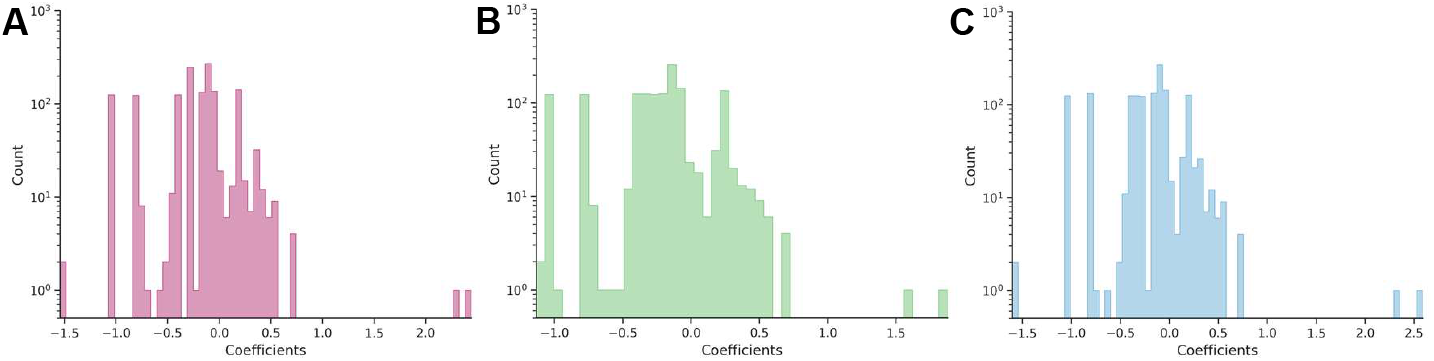
Distribution of third-order epistatic coefficients computed (A) before Power Transform, (B) after the Box-Cox-based Power Transform, and (C) after the Yeo-Johnson-based Power Transform. In both (B) and (C) cases, the Power Transform was applied separately to individual three-dimensional combinatorially complete datasets.

We then applied the Power Transform method, expecting that the distributions of epistatic coefficients would become narrower and that the average epistatic coefficients would decrease in absolute value, as the Power Transform should remove “excessive” nonlinearity, thereby reducing the magnitude of epistatic coefficients.

Application of the Box-Cox-based Power Transform to each hypercube individually confirmed our expectations: the average epistatic coefficient decreased to 0.5432 ± 0.0002 (a 19.4% reduction compared to the reference value) for two-dimensional hypercubes and to 0.454 ± 0.003 (a 1.30% reduction) for three-dimensional hypercubes. Comparison of the pairwise epistasis distributions (Figs. 1A and 1B) showed a statistically significant narrowing (*p* = 6.38⋅10^-9^), whereas comparison of third-order epistatic coefficient distributions (Figs. 2A and 2B) showed no significant effect (*p* = 0.178).

In contrast, application of the Yeo-Johnson-based Power Transform led to a slight increase in epistasis: the average coefficient rose to 0.7040 ± 0.0002 (a 4.45% increase) for two-dimensional hypercubes and to 0.462 ± 0.003 (a 0.43% increase) for three-dimensional hypercubes. The corresponding changes in distributions were not statistically significant: *p* = 0.415 for pairwise coefficients (Fig. 1A and 1C) and *p* = 0.998 for third-order coefficients (Figs. 2A and 2C).

From the distribution of third-order epistatic coefficients, we observe that higher-order epistasis – in this case, third-order – is clearly present both in the original data and after applying the Power Transform (Figs. 2B and 2C), confirming the results reported in (Sailer, Harms, 2017).

### Changes in epistatic coefficients

A narrowing or widening of the entire distribution does not necessarily imply that every coefficient decreased or increased in magnitude, although this is generally expected. To examine this, we calculated the differences in the absolute values of epistatic coefficients for each combinatorially complete dataset. Distributions of these differences are shown in Fig. 3 (for two-dimensional hypercubes) and Fig. 4 (for three-dimensional hypercubes).

**Figure 3.**
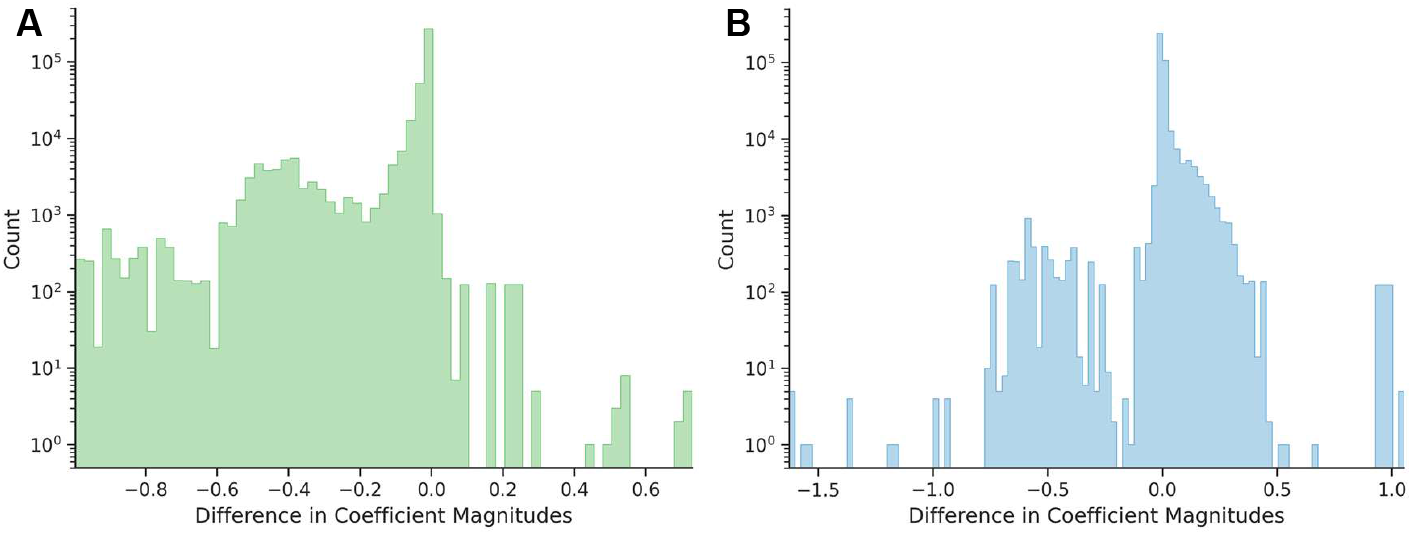
Distribution of changes in absolute values of second-order epistatic coefficients (A) after application of the Box-Cox-based Power Transform, (B) after application of the Yeo-Johnson-based Power Transform.

**Figure 4.**
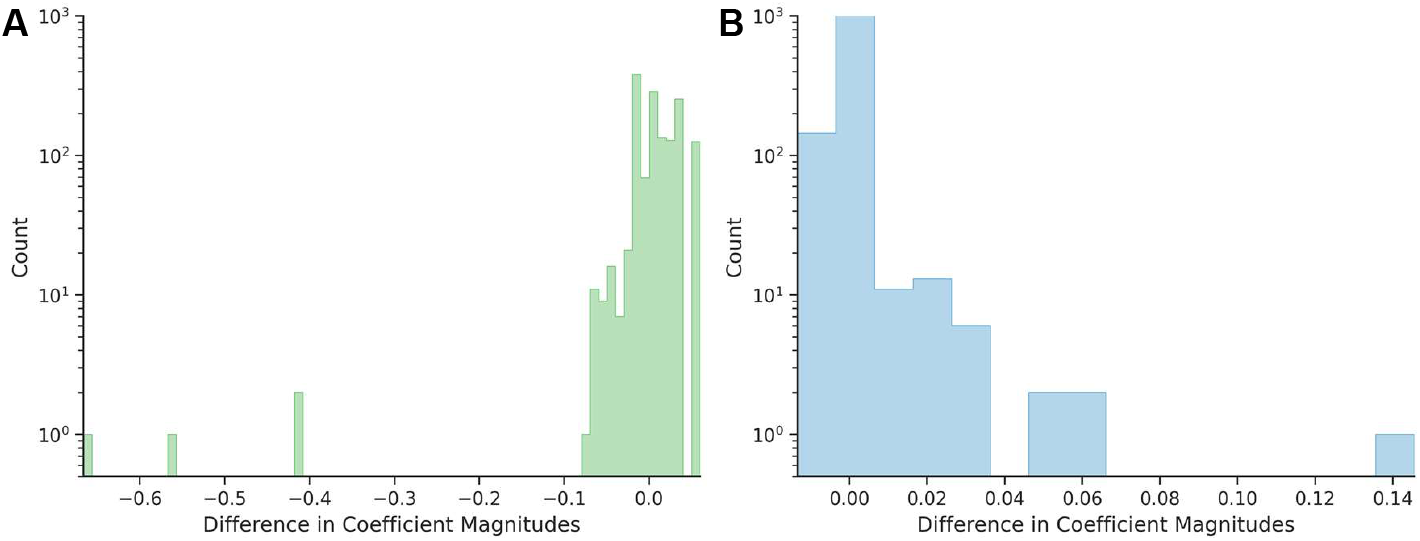
Distribution of changes in absolute values of third-order epistatic coefficients (A) after application of the Box-Cox-based Power Transform, (B) after application of the Yeo-Johnson-based Power Transform.

If the magnitudes of the epistatic coefficients decrease – i.e., move closer to zero – then the differences (post-transform minus pre-transform) should be negative. However, we observe hypercubes, both two-dimensional (Fig. 3) and three-dimensional (Fig. 4), for which the difference is positive, indicating that the absolute values of epistatic coefficients increased after applying the Power Transform. Such cases occur with both the Box-Cox and Yeo-Johnson transformations. A comparison of Fig. 3A and 3B, as well as Fig. 4A and 4B, shows that the Yeo-Johnson method results in significantly higher number of these cases for both two-dimensional and three-dimensional hypercubes.

Overall, for the Box-Cox-based Power Transform, the absolute values of 12,799 second-order coefficients and 941 third-order coefficients increased; for the Yeo-Johnson method, 254,215 second-order coefficients and 650 third-order coefficients increased. Thus, the Power Transform does not always reduce the magnitude of epistatic coefficients. Within our framework, we interpret this as evidence that the Power Transform does not reduce nonlinearity for some hypercubes.

### Application of Power Transform to the whole landscape

Beyond applying the Power Transform to individual hypercubes, it is desirable to apply it to entire landscapes. However, when attempting this on the GFP landscape, we found that, for landscapes generated by random mutagenesis, the Power Transform generally fails. It appears that, for the Power Transform to function, the landscape must satisfy the following condition: for any single mutation present in the dataset, there must exist at least one pair of genotypes that differ only at that site while being identical at all others.

It can be shown that this condition is satisfied if the landscape represents a connected component – specifically, if for every genotype in the component, there exists at least one other genotype at Hamming distance 1.

We partitioned the GFP landscape into connected components as described in Materials and Methods. We found that the experimental GFP landscape contains only one large connected component, consisting of 20,872 genotypes. We applied the Yeo-Johnson-based Power Transform to this component and computed epistatic coefficients for all embedded hypercubes (two- and three-dimensional), but using phenotype values linearized accross the entire component. The distributions of epistatic coefficients and the differences in their absolute values, are presented in Fig. 5.

**Figure 5.**
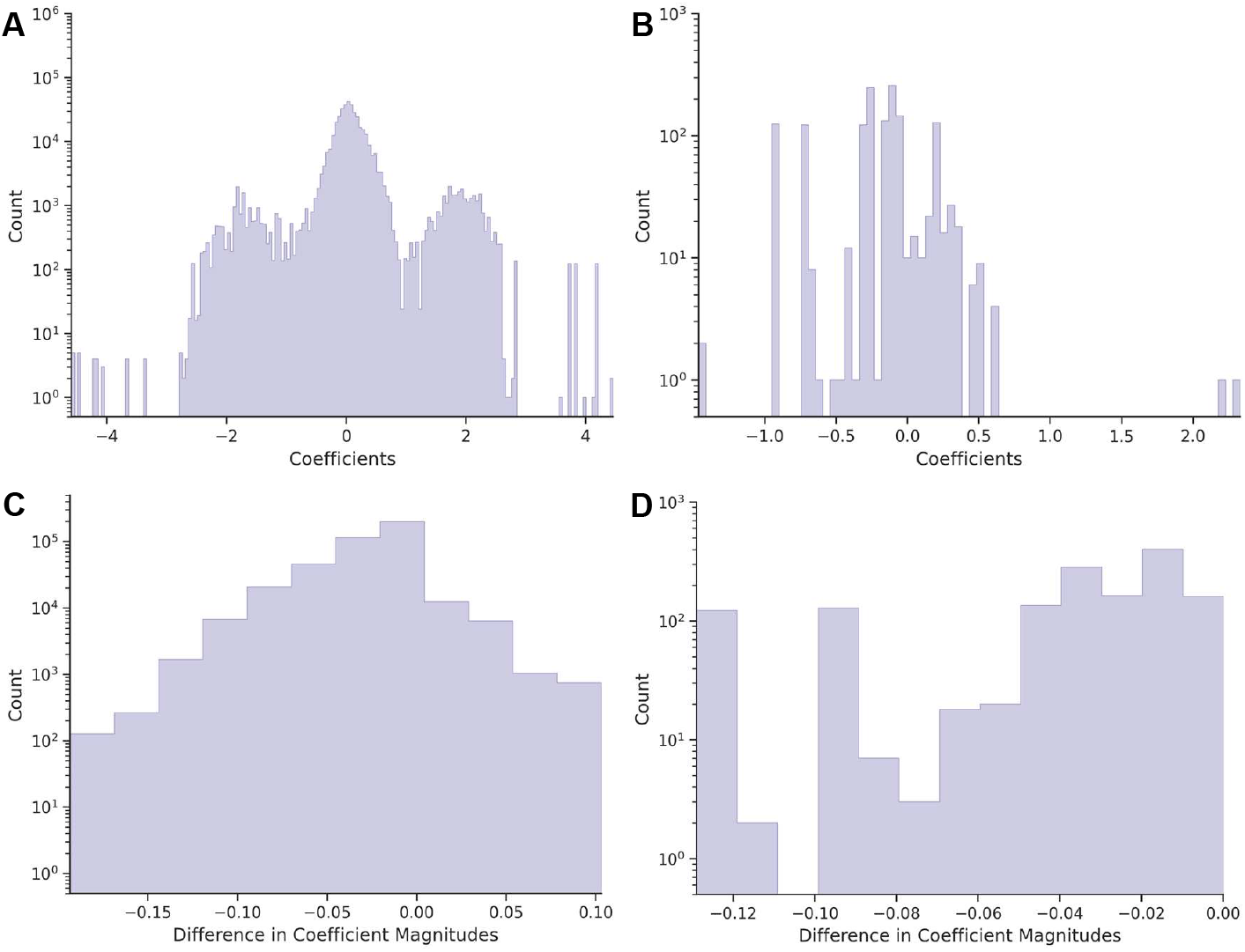
Distribution of epistatic coefficients (A) of the second order, (B) of the third order, and changes in absolute values of coefficients (C) of the second order, (D) of the third order after application of Yeo-Johnson-based Power Transform across the entire largest connected component of the GFP landscape.

The average epistasis for two- and three-dimensional hypercubes was 0.648 and 0.152, respectively – corresponds to reductions of 3.86% and 11.6% compared to initial reference values.

As shown in Figs. 5C and 5D, the majority of two-dimensional epistatic coefficients (note the logarithmic scale on the vertical axis) and all three-dimensional decreased in magnitude after applying the Yeo-Johnson-based Power Transform to the largest connected component. However, in a subset of two-dimensional cases, the absolute value of the epistatic coefficient increased: this occurred in 6.3% of hypercubes (25,491 out of 403,162).

## Discussion

In this work, we introduced and validated a conceptual framework for applying the Power Transform method to large fitness landscapes, including those generated by (quasi-)random mutagenesis. The approach involves partitioning the landscape into connected components and applying a Yeo-Johnson-based Power Transform to each component individually. While the Box-Cox-based Power Transform was found to be more effective on individual hypercubes, it is less universally applicable because it cannot accommodate non-positive values – not only in the observed phenotypes but also in their linearized counterparts. In large landscapes, such as the well-studied GFP landscape (which spans up to 15 mutational steps), non-positive linearized values are highly probable, rendering Box-Cox version of Power Transform inapplicable.

We demonstrated our concept on the canonical GFP landscape obtained via random mutagenesis (Sarkisyan *et al*., 2016). The only large connected sublandscape amenable to Power Transform comprised 20,872 genotypes. After applying the Yeo-Johnson-based Power Transform, pairwise epistasis decreased by 3.86%, and third-order epistasis by 11.6%.

The original GFP study reported a sigmoidal relationship between the logarithm of fluorescence and a proxy for fitness potential – namely, protein stability (see Fig. 5 of (Sarkisyan *et al*., 2016)). This nonlinearity arises because two mildly destabilizing mutations, when combined, may push the protein beyond its stability threshold, preventing proper folding, chromophore formation, and thus, being fluorescent. This sigmoidal behavior was also captured by a neural network. The trimodal distribution of pairwise epistatic coefficients (Fig. 1A) is a direct consequence of this sigmoidal landscape, reflecting three distinct scenarios:

1. Distinct negative epistasis (left peak, ≈ –2): A stable reference genotype (00) and its single mutants (01, 10) fluoresce, so the double mutant (11) is expected to fluoresce as well. If, however, the double mutant falls beyond the sigmoidal stability threshold and fails to fluoresce, the observed-expected difference yields a distinct negative epistatic coefficient.
2. Distinct positive epistasis (right peak, ≈ +2): A destabilized reference genotype (00) and its single mutants (01, 10) do not fluoresce, so the double mutant (11) is expected to remain dark. If stabilizing mutations shift the double mutant 11 left of the stability threshold, enabling fluorescence, the epistatic coefficient becomes definitely positive.
3. Near-zero epistasis (central peak): It occurs when fluorescence changes (nearly) expectedly. For example, when all four genotypes (00, 01, 10, 11) either fluoresce or are dark, or when 00 and 01 are bright while 10 and 11 are dark, yielding negligible epistasis.

Comparing Box-Cox and Yeo-Johnson transformations on individual hypercubes from GFP data reveals a counterintuitive result: Box-Cox achieves better linearization than Yeo-Johnson, despite the latter’s ability to handle both positive and non-positive values and its broader functional scope. One might expect Yeo-Johnson transformation – as more generally applicable – to perform equally well or better than Box-Cox version. However, we did not investigate why Box-Cox outperforms Yeo-Johnson in local contexts of individual hypercubes.

This observation might hint to a critical distinction: optimizing the Power Transform parameters does not necessarily maximize linearization. In other words, “best Power Transform” and “best linearization” are not synonymous. Indeed, for both methods, we observed cases where the transform increased epistatic coefficients, further indicating that Power Transform application does not guarantee reduced nonlinearity.

Surprisingly, Yeo-Johnson-based Power Transform reduced epistasis across the full landscape but, on average, increased it within individual hypercubes. We speculate that the global transform finds compromise parameters that effectively suppresses landscape-wide nonlinearity, whereas local hypercubes lack sufficient structure or signal for the same parameters to yield linearization. However, we did not probe this mechanism deeply; we note it as an observation worthy of further study.

In summary, we have developed and demonstrated a viable concept: Power Transform methods – specifically, based on Yeo-Johnson transformation – can be applied to large fitness landscapes, specifically obtained by (quasi-)random mutagenesis. The broader implications and future utility of this approach remain to be explored.

